# Translation of monosynaptic circuits underlying amygdala fMRI neurofeedback training

**DOI:** 10.1101/2024.03.15.585281

**Authors:** Lucas Trambaiolli, Chiara Maffei, Evan Dann, Claudinei Biazoli, Gleb Bezgin, Anastasia Yendiki, Suzanne Haber

**Author notes:** **Corresponding authors:** Lucas Trambaiolli, Suzanne Haber.

## Abstract

**Background:** fMRI neurofeedback targeting the amygdala is a promising therapeutical tool in psychiatry. It induces resting-state functional connectivity (rsFC) changes between the amygdala and regions of the salience and default mode networks (SN and DMN, respectively). We hypothesize these rsFC changes happen on the amygdala’s underlying anatomical circuits.

**Methods:** We used the coordinates from regions of interest (ROIs) from studies showing pre-to-post-neurofeedback changes in rsFC with the left amygdala. Using a cross-species brain parcellation, we identified the homologous locations in non-human primates. We injected bidirectional tracers in the amygdala of adult macaques and used bright- and dark-field microscopy to identify cells and axon terminals in each ROI. We also performed additional injections in specific ROIs to validate the results following amygdala injections and delineate potential disynaptic pathways. Finally, we used high-resolution diffusion MRI data from four *post-mortem* macaque brains and one *in vivo* human brain to translate our findings to the neuroimaging domain.

**Results:** The amygdala had significant monosynaptic connections with all the SN and DMN ipsilateral ROIs. Amygdala connections with the DMN contralateral ROIs are disynaptic through the hippocampus and parahippocampal gyrus. Diffusion MRI in both species benefitted from using the ground-truth tracer data to validate its findings, as we identified false-negative ipsilateral and false-positive contralateral connectivity results.

**Conclusions:** Amygdala neurofeedback modulates the SN and DMN through monosynaptic connections and disynaptic pathways - including hippocampal structures involved in the neurofeedback task. Neurofeedback may be a tool for rapid modulation and reinforcement of these anatomical connections, leading to clinical improvement.

## Introduction

Functional Magnetic Resonance Imaging (fMRI) neurofeedback modulates specific brain regions in real time through self-elicited mental strategies [1]. It is considered a potential therapeutic approach in psychiatry for several disorders, including depression, anxiety, and substance abuse [2–4]. Comparing resting-state functional connectivity (rsFC) before and after neurofeedback provides insights into which brain networks support the long-lasting effects of neuromodulation [5, 6]. However, fMRI is an indirect method for connectivity analysis. Delineating the hard-wired, monosynaptic connections that underlie these rsFC results will lead to: (i) identifying the circuitries underlying neurofeedback mechanisms; (ii) probing those circuits with animal models; (iii) developing potential biomarkers; (iv) guiding personalized neurofeedback protocols. This study aims to determine the extent to which rsFC changes elicited by neurofeedback modulation represent changes in anatomic, monosynaptic connections. We use NHP anatomic tracing experiments and high-resolution diffusion MRI (dMRI) data in macaques and humans to determine the most likely monosynaptic connectivity changes following neurofeedback intervention.

We focused on connectivity changes following fMRI neurofeedback of the amygdala, a successful neurofeedback target [7, 8]. The current hypothesis is that clinical improvement following amygdala neurofeedback results from its modulation of two large-scale networks: the salience and the default mode networks (SN and DMN, respectively) [8–12]. Although connections between the amygdala and SN and DMN nodes have been described [13–34], these are large regions with several subdivisions, each with different connectivity patterns [15, 23, 24, 27, 29, 30]. Here, we combined multi-modal multi-species data to delineate the anatomical circuits connecting the amygdala and the specific DMN and SN sublocations (or regions-of-interest - ROIs) modulated by amygdala neurofeedback [5, 12, 35]. Specifically, we: *1.* Identified the equivalent ROIs in the non-human primate (NHP) brain. *2.* Analyzed the anatomical connections between each sublocation and the amygdala. *3.* Tested whether the same connections could be identified using submillimeter *ex vivo* dMRI tractography data in NHP. *4.* Tested whether these connections could also be identified in human submillimeter *in vivo* dMRI.

Our results show that the main rsFC changes following neurofeedback are likely sustained by monosynaptic connections between the amygdala and ipsilateral ROI sublocations of the SN and DMN nodes. Amygdala connections with contralateral ROIs are disynaptic, likely through hippocampal and parahippocampal gyrus (PHG) regions involved in neurofeedback tasks [11, 36, 37]. This anatomical delineation allows for future neurofeedback studies probing those circuits and related mechanisms in human and animal models and using neurofeedback to identify biomarkers or personalized targets in clinical samples.

## Methods

### Step 1: Translating rsFC ROIs from human to the NHP brain

We extracted the rsFC ROIs from all three studies [5, 12, 35] that used seed-based connectivity analysis to evaluate an fMRI neurofeedback protocol based on positive autobiographical memory recall to up-regulate the BOLD signal of the left amygdala. There were 16 ROIs (Table 1) in which rsFC with the left amygdala changed after neurofeedback, including some, but not all, ROIs of the SN (dorsal anterior cingulate cortex - dACC, anterior insula – AI, and lateral prefrontal cortex - LPFC) and DMN (middle frontal gyrus - MFG, temporal pole - TP, hippocampus, parahippocampal gyrus – PHG, precuneus, posterior cingulate cortex – PCC and thalamus). The coordinates of these ROIs were transformed to the MNI space based on Lancaster, et al. [38].

**Table 1.** List of center or peak coordinates of nodes showing amygdala-rsFC changes when comparing pre- and post-amygdala neurofeedback training in human subjects and the equivalent coordinates in the homologous structures of the macaque brain. Nodes are grouped as part of the human Salience or Default Mode Networks. Coordinates in the human brain are reported in the MNI brain template (center), and coordinates in the macaque brain in the F99 brain template. * = coordinates estimated based on figures.

| Original ROI label | Human coordinates<br>(MNI space) |  |  | NHP coordinates<br>(F99 space) |  |  |
| --- | --- | --- | --- | --- | --- | --- |
|  | x | y | z | x | y | z |
| **Ipsilateral (Left Hemisphere) Salience Network nodes** |  |  |  |  |  |  |
| Dorsal anterior cingulate cortex (dACC) | -2 | 25 | 30 | -2 | 10 | 15 |
| Lateral prefrontal cortex (LPFC) | -50 | 14 | 0 | -24 | 7 | 0 |
| Anterior Insula (AI) | -37 | 21 | 5 | -21 | 3 | 1 |
| **Ipsilateral (Left Hemisphere) Default Mode Network nodes** |  |  |  |  |  |  |
| Middle frontal gyrus | -30 | 15 | 44 | -13 | 11 | 17 |
| Temporal pole | -43 | 3 | -24 | -18 | 3 | -6 |
| Parahippocampal gyrus (PHG) | -26 | -4 | -19 | -8 | -3 | -14 |
| Angular gyrus* | -44 | -53 | 28 | -21 | -23 | 15 |
| Medial precuneus* | -7 | -59 | 40 | -1 | -24 | 20 |
| Lateral precuneus | -27 | -54 | 46 | -14 | -23 | 19 |
| **Contralateral (Right Hemisphere) Default Mode Network nodes** |  |  |  |  |  |  |
| Middle frontal gyrus | 28 | 45 | 33 | 17 | 14 | 14 |
| Temporal pole | 45 | 18 | -29 | 22 | 6 | -9 |
| Parahippocampal gyrus (PHG) | 30 | -27 | -23 | 17 | -5 | -18 |
| Hippocampus | 30 | -26 | -2 | 15 | -12 | -8 |
| Posterior cingulate cortex | 3 | -47 | 31 | 2 | -23 | 11 |
| Medial precuneus | 13 | -31 | 50 | 6 | -12 | 17 |
| Thalamus | 17 | -24 | -2 | 9 | -14 | -1 |

We used the “Regional Map” parcellation, a standard for cross-species comparisons [39], to identify equivalent ROIs across NHPs and humans. Details about this parcellation are included in the *Supplementary Information*. The equivalent ROIs in the macaque brain were manually placed according to homologous parcels and anatomical landmarks. Importantly, in this study, we used human terminology when referring to the NHP ROIs (e.g., although we list ROIs in the “angular gyrus” and “middle frontal gyrus” macaques don’t have these gyri in the strict sense).

### Step 2: Identification of anatomical connections using NHP tract-tracer data

Our laboratory has an extensive collection of bidirectional tracer injections placed throughout cortical and subcortical areas of adult male macaque monkeys. From this database, we selected 12 injection sites (four placed in *Macaca mulatta*, four in *Macaca fascicularis*, and four in *Macaca nemestrina*). For the seven injections in the amygdala (Figure 1A), we evaluated connections with each SN and DMN ROIs from Table 1. We used the additional five injections in specific ROIs (anterior insula, lateral precuneus, hippocampus, posterior cingulate cortex, and thalamus) to validate the connectivity patterns with the amygdala. The surgical and histological procedures are detailed in the *Supplementary Information*.

**Figure 1.**
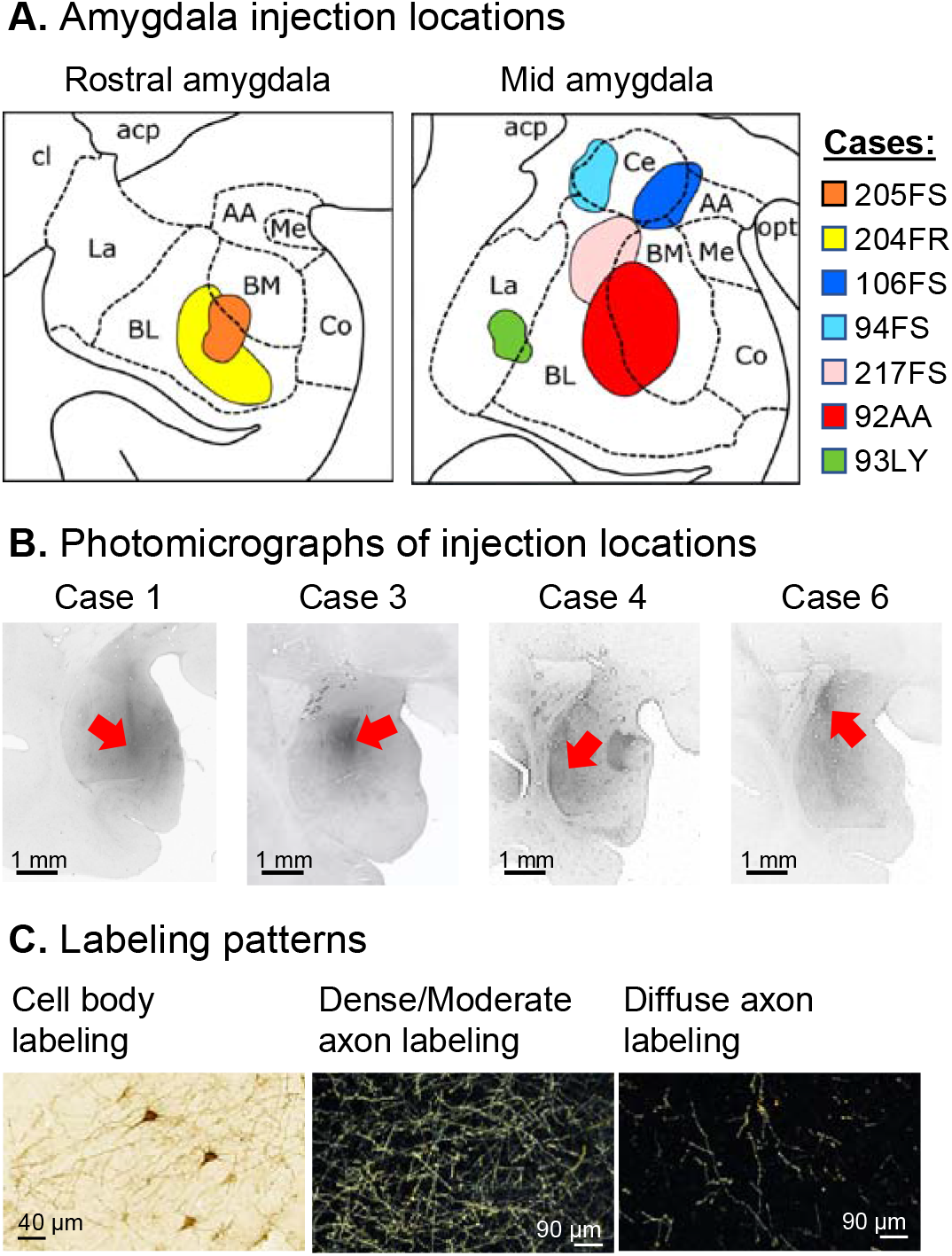
Amygdala Injection sites and labeling patterns. **A.** Schematic of injection sites at approximately the same rostro-caudal level in the macaque amygdala. Dotted lines = nuclei borders. Colored areas = individual cases. **B.** Coronal sections of the macaque amygdala showing different injection locations. Scale bar, 1 mm. **C.** Examples of dense/moderate and diffuse terminal fields. *Abbreviations*: AA = Anterior area; acp = anterior commissure – posterior limb; BL = Basolateral nucleus; BM = Basomedial nucleus; Ce = Central nucleus; cl = claustrum; Co = Cortical nucleus; La = Lateral nucleus; Me = Medial nucleus; opt = optic tract.

Using StereoInvestigator software (MicroBrightField Bioscience, U.S.A), we charted the retrogradely labeled cells in the ROIs under light-field microscopy at 20 x (Figure 1B, left) [40–42]. We used dark-field microscopy under 1.6 x, 4 x, and 10 x objectives with Neurolucida software (MicroBrightField) to outline dense or light axon terminal projections in the ROIs. We labeled condensed groups of fibers visible at 1.6 x with discernible boundaries as ‘dense projections’ (Figure 1B, center), and groups of fibers where individual terminals could be discerned as ‘light projections’ (Figure 1B, right) [40, 42].

### Step 3: Identification of anatomical connections using NHP tractography data

The NHP postmortem submillimeter dMRI data was collected from four adult animals, with a total scan time of 47 hours per brain (MRI acquisition details can be found in the *Supplementary Information*). The dMRI data underwent a preprocessing pipeline that included denoising [43] and correction for Gibbs ringing [44], signal drift [45], eddy-currents [46], and bias fields [47]. We fit fiber orientation distribution functions (fODF) to the pre-processed data using multi-shell multi-tissue constrained spherical deconvolution (MSMT-CSD [48]) in MRtrix3 [47]. The D99 macaque atlas [49] was transformed to each individual brain after registering the D99 magnetization transfer ratio (MTR) template volume to the individual b=0 volume using the robust affine registration in FreeSurfer (mri_robust_register [50]). The left amygdala was extracted from the D99 atlas, binarized, and dilated by 2 voxels in MRtrix3 to include the surrounding white matter. We performed probabilistic tractography in MRItrix3 seeding in every voxel within this mask (350 seeds per voxel). The following tractography parameters were used: step-size = 0.25 mm, maximum angle threshold = 30°, fODF peak threshold = 0.06, and maximum length = 150 mm.

The location of each rsFC ROI was identified as a single point on the D99 macaque brain based on anatomical landmarks. These point coordinates were mapped to each individual brain using the transforms from the registration described above. For each point, we found its nearest point along the white-gray matter boundary. Spherical ROIs were defined with a 1.5 mm radius around these points. Streamlines connecting the left amygdala and each of the ROIs included in this analysis were manually dissected using Trackvis (v.0.6.1; http://www.trackvis.org). Streamlines connecting the left amygdala with each ROI were filtered to only include those streamlines ending or originating inside the amygdala mask.

### Step 4: Human tractography analysis

We used submillimeter-resolution dMRI data from a publicly available and pre-processed dataset [51] (see details in the *Supplementary Information*). Processing followed similar steps to those previously described for the NHP data. We fit fODFs to the preprocessed dMRI data using multi-shell multi-tissue constrained spherical deconvolution (MSMT-CSD) in MRtrix3. Cortical parcellations and subcortical segmentations were obtained from the T1 data using FreeSurfer [52–54]. The left amygdala was extracted from the segmentation, binarized, and inflated by 1 voxel. We performed probabilistic tractography in MRtrix3, seeding in every voxel within the amygdala (100 seeds per voxel). The following tractography parameters were used: step-size = 0.38 mm, maximum angle threshold = 45°, maximum length = 150 mm. We mapped the ROI coordinates from the MNI space to the individual space using the registration procedures described in the NHP analysis.

Streamlines connecting the left amygdala and each ROI from Table 1 were manually dissected using Trackvis. For the amygdala, a sphere was created around the center coordinates extracted from seed-based rsFC studies [5, 12]. A 10 mm radius was used to include the surrounding white matter. For all other cortical ROIs, spheres of 7 mm radius with their centers at the border between white and gray matter closest to the ROI coordinates.

## Results

### Step 1: Cross-species ROIs selected for this study

Based on anatomical landmarks, we identified the equivalent 16 ROIs in the NHP brain, and the resulting center coordinates in the F99 space are listed in Table 1. The SN ROIs in the NHP brain included the ipsilateral (left hemisphere) dACC (at the genu of the corpus callosum including area 24, Figure 2A-left), AI (at the rostral portion of the circular sulcus including area AI, Figure 2B-left), and LPFC (caudal area 47/12 extending to ProM at the dorsal lip of the rostral part of lateral fissure Figure 2C-left).

**Figure 2.**
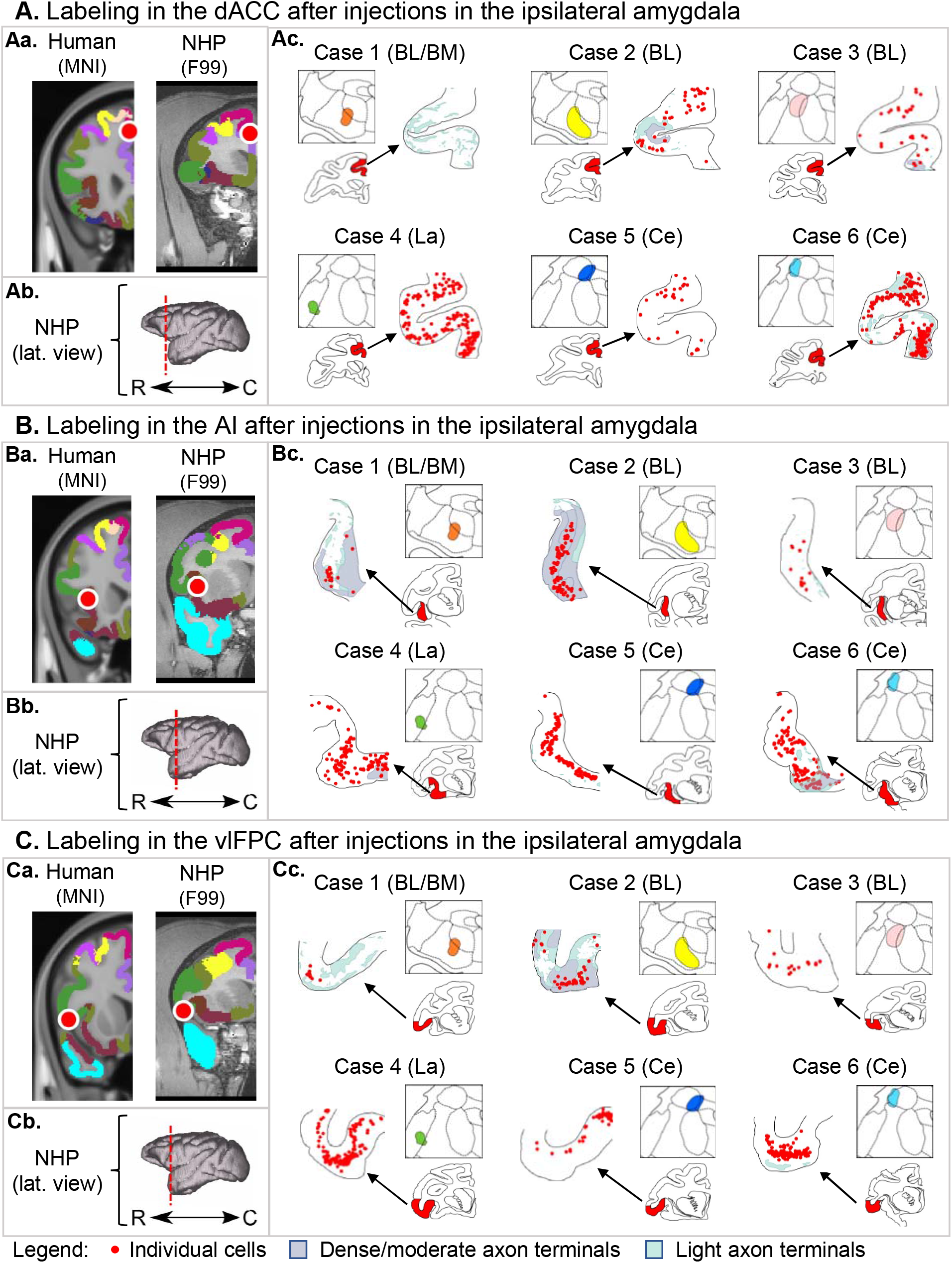
Amygdala connections with the ipsilateral SN nodes. **Aa.** Red circles indicate the peak location of the rsFC changes after amygdala neurofeedback for the dACC in the human MNI template and the homologous location in the macaque F99 template. **Ab.** 3D models represent the rostro-caudal location of coronal slices from each node. **Ac.** For each case, the respective injection is shown in the square box, and the schematic coronal sections highlight in red the location with connectivity chartings. Individual cells are shown as red dots, dense/moderate terminals as light blue shaded areas, and diffuse terminals as light green shaded areas. The same organization is followed for ROIs in the AI (**B**) and vlPFC (**C**). *Abbreviations:* BL = basolateral nucleus, BM = basomedial nucleus, C = caudal, Ce = central nucleus, La = lateral nucleus, R = rostral.

The ipsilateral ROIs of the DMN included the MFG (at the ventral bank of the superior arcuate sulcus, in the border of areas 8AB, 8B, and 9/46D, Figure 3A-left) in the frontal cortex. In the temporal cortex, the TP (at the ventral bank of the circular sulcus including areas IPro and TPPro Figure 3B-left) and PHG (dorsal to the rhinal fissure, at the border of areas EI, ELC, and ER, Figure 3C-left). Finally, in the parietal cortex, ROIs included the Lateral Precuneus (at the lip of the ventral bank of the intraparietal sulcus, including areas POaE/LIPE and PG, Figure 3D-left), Medial Precuneus (at the ventral bank of the cingulate sulcus, including areas PGm and 31, Figure 3E-left), and Angular Gyrus (AG, in the lateral fissure, including the border of areas PGOp/ReI, PaAC and Tpt, Figure 3F-left).

**Figure 3.**
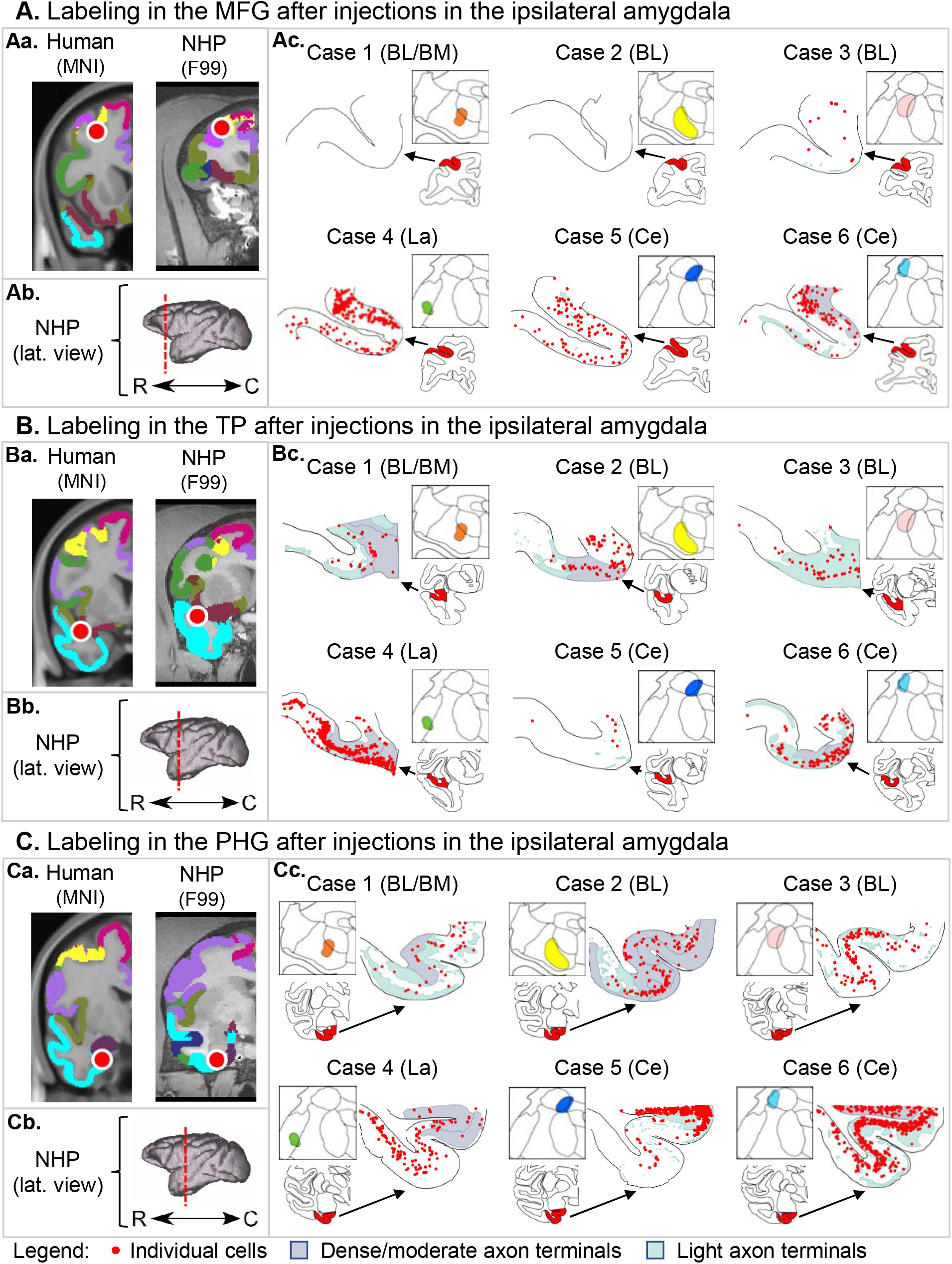
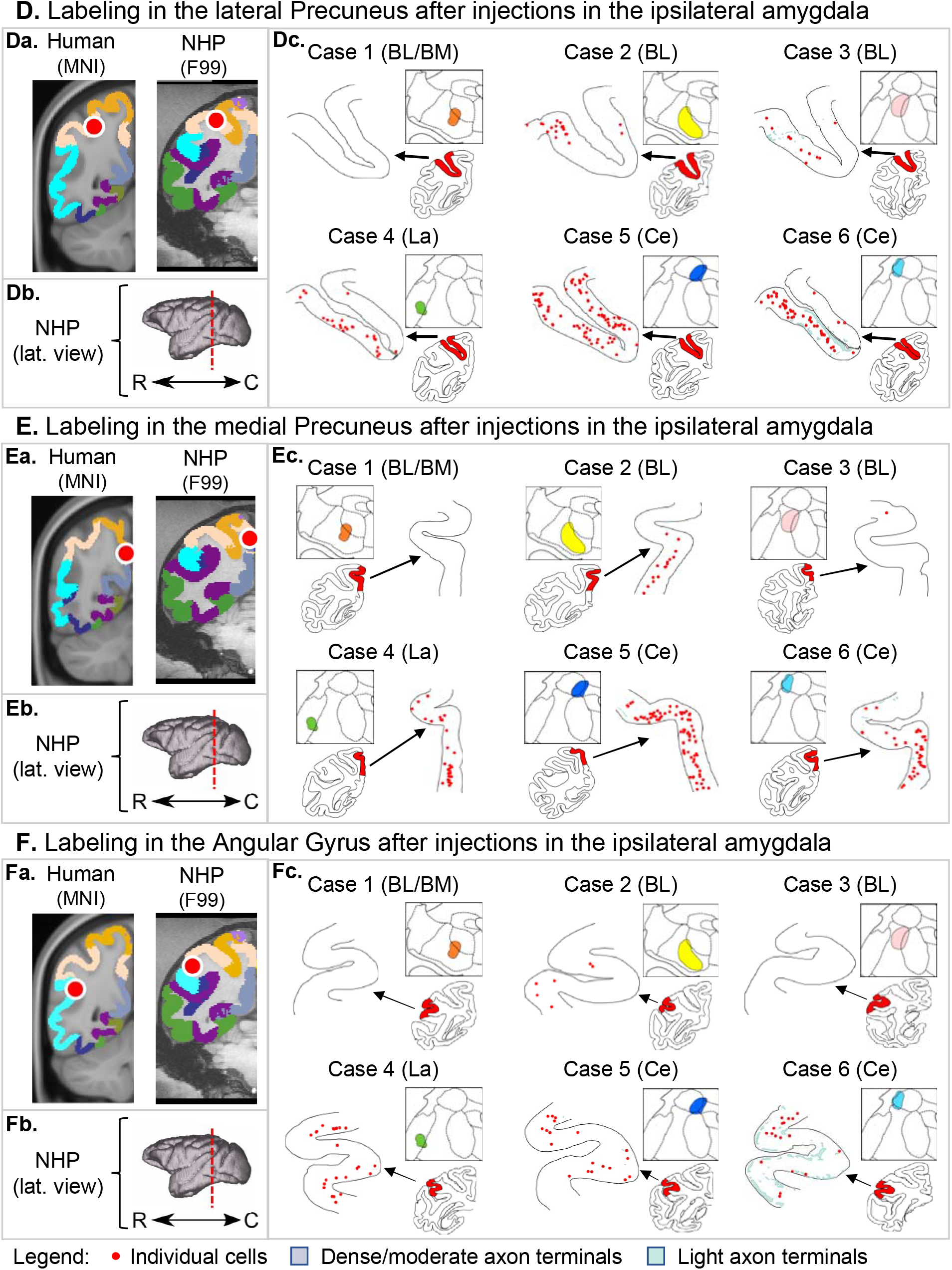
Amygdala connections with the ipsilateral DMN nodes. **Aa.** Red circles indicate the peak location of the rsFC changes after amygdala neurofeedback for the Middle Frontal Gyrus in the human MNI template and the homologous location in the macaque F99 template. **Ab.** 3D models represent the rostro-caudal location of coronal slices from each node. **Ac.** For each case, the injections in the amygdala are shown in the square box, and the schematic coronal sections highlight the location with connectivity chartings. Individual cells are shown as red dots, dense/moderate terminals as light blue shaded areas, and diffuse terminals as light green shaded areas. The same organization is followed for ROIs in the Temporal Pole (**B**), Parahippocampal Gyrus (**C**), Lateral Precuneus (**D**), Medial Precuneus (**E**), and Angular Gyrus (**F**)*Abbreviations:* BL = basolateral nucleus, BM = basomedial nucleus, C = caudal, Ce = central nucleus, La = lateral nucleus, R = rostral.

The DMN ROIs in the contralateral hemisphere included the MFG (at the lower bank of the principal sulcus, including areas 9/46V and 46V) in the frontal cortex. In the temporal cortex, TP (at the dorsolateral portion of the anterior temporal lobe, at area TPPro extending to ST1 - Fig 6A-center) and PHG (at the lateral bank of the rhinal fissure, including areas TLR/R36 and 35 - Fig 6A-right). In the parietal cortex, PCC (area 23 in the cingulate gyrus – Fig 6B) and Medial Precuneus (dorsal bank of the cingulate sulcus, in area 31 extending to areas 23 and 3). And subcortical ROIs include the dorsal Hippocampus (Fig 6A-left) and Thalamus (at the transition between ventral, centromedial, mediodorsal, and pulvinar nuclei – Fig 6C).

### Step 2: Anatomical connections identified using NHP tract-tracer data

Bidirectional tracer injections in the amygdala showed monosynaptic connections with the ipsilateral dACC, AI, and LPFC ROIs within the SN Figure 2). Importantly, the basolateral (BL), lateral (La), and lateral central (Ce, case 6) amygdala nuclei, had bidirectional connections with these three ROIs. An anterograde BL injection (Case 7, not shown) was consistent with the spatial patterns of axon terminals observed in the other BL injections. To validate the existence and specificity of the observed connections, we identified a small bidirectional tracer injection in the AI (Figure 4B) that showed bidirectional connectivity patterns spread along all amygdala nuclei, consistent with the results in Figure 2B.

**Figure 4.**
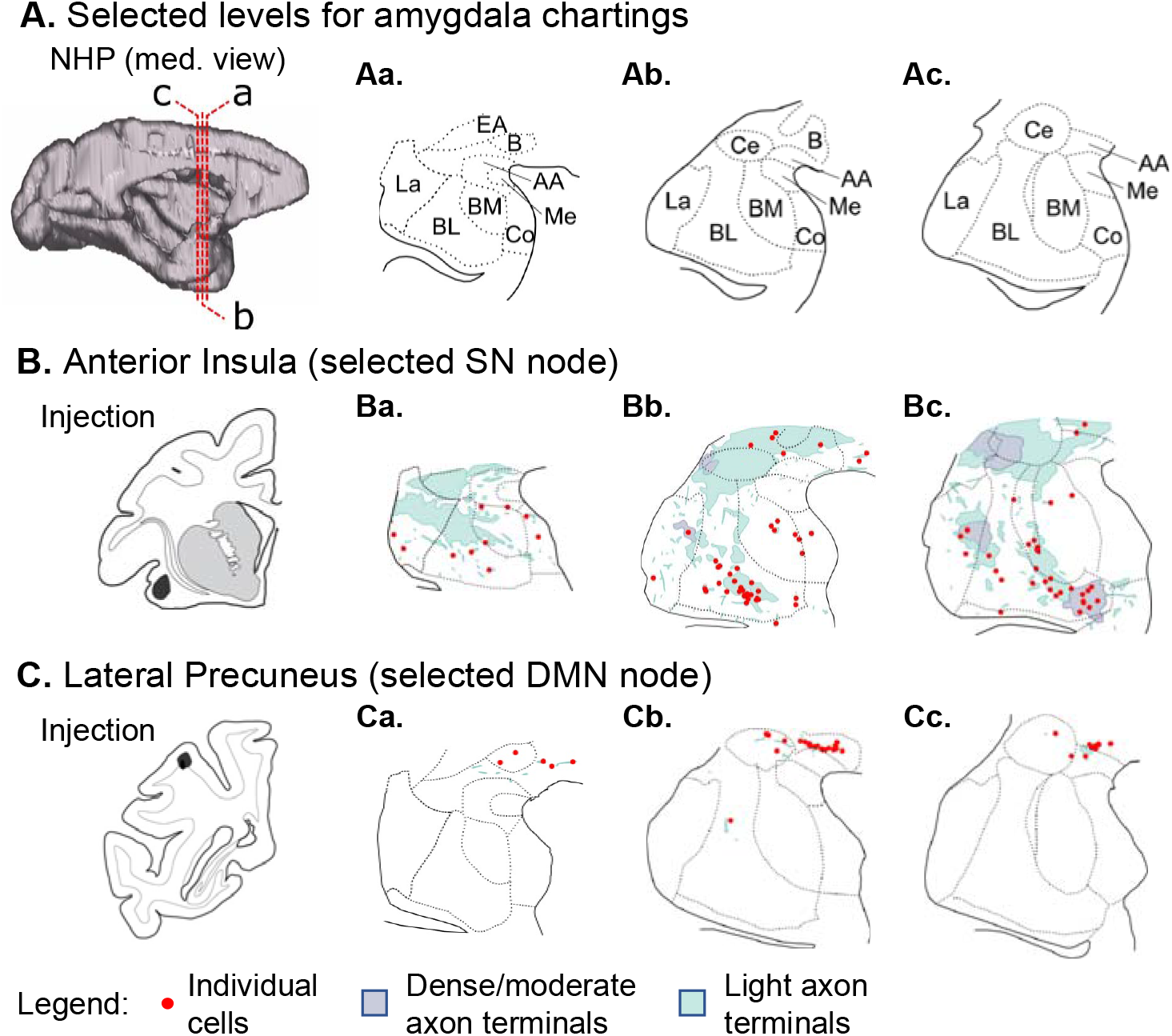
Validation of amygdala connections after cortical injections. **A.** 3D representation of the three rostro-caudal levels (a-b) used in the chart cells and terminals in the amygdala, and the respective coronal slices with cytoarchitectonic divisions based on the Paxinos atlas. Labeling of cells (red dots), and dense/moderate (light blue) and diffuse (light green) terminal fields in the amygdala after bidirectional tracer injections in regions homologous to the Anterior Insula (**B**), and Lateral Precuneus (**C**) regions with resting-state functional connectivity changes after amygdala neurofeedback. *Abbreviations:* AA = anterior amygdaloid area. BL = basolateral nucleus, BM = basomedial nucleus, C = caudal, Ce = central nucleus, La = lateral nucleus, R = rostral.

The amygdala was also anatomically interconnected with the ipsilateral DMN sublocations modulated by neurofeedback (Figure 3). ROIs closer to the amygdala (TP and PHG, Figure 3B-C) were bidirectionally connected with all injection locations. The MFG (Figure 3A), precuneus (Figure 3D-E), and angular gyrus (Figure 3F) showed dense labeling with La and Ce injection sites but scarce labeling with BL injections (Cases 2 and 3). None of these DMN ROIs connected with the injection in BL/BM (Case 1). A validation injection in the lateral precuneus (Figure 4C) showed concentrated labeling in the dorsal bank of the amygdala, including the Basal and Ce nuclei. Although this injection is lateral to the original ROI, these results are partially consistent with those observed in the amygdala injections (Figure 3D), except for the lack of labeling in the La nucleus. Importantly, labeling in parietal structures (precuneus and angular gyrus) after injections in the amygdala and labeling in the amygdala after injection in the precuneus showed predominantly retrograde labeling.

Our tracing data showed sparse monosynaptic connections from the left amygdala to the contralateral hemisphere and no connections with the specific contralateral DMN ROIs. Importantly, amygdala neurofeedback is also associated with changes in the hippocampus and parahippocampal gyrus (PHG) [11, 12, 36, 37, 55], regions also anatomically interconnected with the amygdala [14, 26, 28]. Thus, we evaluated if the amygdala is connected with the contralateral DMN ROIs through the hippocampus and PHG. Supplementary Figure 1B shows anatomical labeling in the left hippocampus and PHG for injections in the left amygdala. Briefly, all cases presented labeling in the amygdalohippocampal area. BL and La injections showed labeling in rostral CA1’ and CA3 subfields and dense labeling in the transition between the subiculum and prosubiculum fields, extending to areas 35, 36, and TF in the PHG. A similar but weaker pattern is also observed in Ce (Case 5). A validation injection extending from CA1, ProS, and part of the Subiculum in the hippocampus to areas 35, 36, and TF in the PHG (Supplementary Figure 1C) showed spatial labeling in the amygdala consistent with those observed in Supplementary Figure 1B.

Supplementary Figure 2 illustrates connections between the left hippocampus and PHG and the contralateral DMN ROIs (right hemisphere). Shortly, the injection in the hippocampus and PHG showed anatomical connections with the contralateral hippocampus, PHG, and TP nodes of the DMN (Supplementary Figure 2 A). However, this injection did not include all structures in the hippocampus and PHG. To evaluate if the remaining contralateral DMN nodes listed in Table 1 connected with other hippocampal and PHG subnuclei, we placed two additional injections in two of these nodes: the right PCC (Supplementary Figure 2B) and right thalamus (Supplementary Figure 2C). We observed axon terminal and cell labeling in the left hippocampus and PHG, respectively.

### Step 3: Anatomical connections identified with NHP tractography

Using submillimeter dMRI tractography from four animals, we could correctly identify structural connections between the left amygdala and all ipsilateral ROIs. Figure 5 shows examples of well-defined tracts connecting the left amygdala and all ipsilateral ROIs of the SN and DMN in one animal. Additional connections and the results from the other animals are shown in Supplementary Figures 3-5. Importantly, inconsistent with the tracer data, connections between the amygdala and the MFG and LPFC were among those with the fewest streamlines compared to other connections.

**Figure 5.**
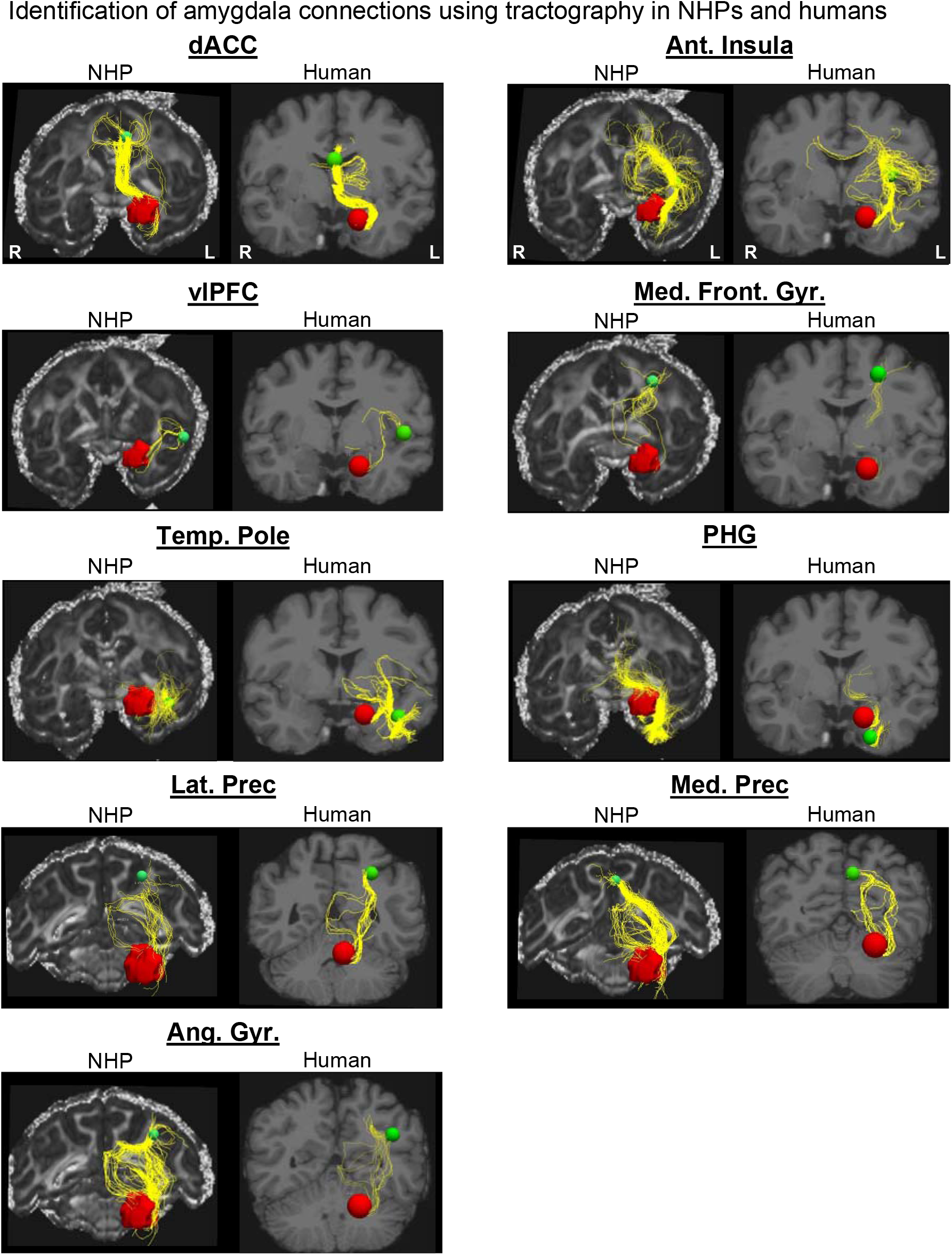
Identification of amygdala connections using dMRI tractography in NHPs and humans. Reconstruction of streamlines (yellow) connecting the amygdala (red) with all ipsilateral nodes (green) within the SN and DMN. For each node, results from one NHP brain are shown on the left, and results from the human brain on the right.

Tractography data also showed streamlines connecting the left amygdala with several contralateral ROIs, disagreeing with the tracer data. We compared the tractography and the tracer data at different locations along these tracts to identify where tractography errors occurred. Supplementary Figure 6A shows two sets of streamlines erroneously connecting the left amygdala and the right medial precuneus in one representative case. After leaving the amygdala, the anterior streamlines follow the same direction as the amygdalofugal fibers (Supplementary Figure 6B-C). However, posteriorly, these streamlines enter the fornix (Supplementary Figure 6D), which is inconsistent with the results from the tracer data. The posterior false positive connection follows the same trajectory as the stria terminalis observed in the tracer data (Supplementary Figure 6E-G). However, similar to the anterior false positive, streamlines erroneously follow through the fornix to the contralateral hemisphere.

### Step 4: Anatomical connections identified with human tractography

Using submillimeter human dMRI tractography, we successfully identified connections between the left amygdala and all ipsilateral ROIs. Amygdala connections with regions such as the dACC, AI, PHG, TP, and Precuneus showed the same clear tracts with dense concentrations of streamlines (Figure 5) as observed in the NHP dMRI tractography data. Consistent with the NHP dMRI tractography data, amygdala-LPFC, and amygdala-MFG connections presented fewer, sparser streamlines than other connections. Similar to tractography results in NHP, false positive connections were also identified connecting the left amygdala with contralateral ROIs (e.g., streamlines traveling contralaterally through the fornix, Supplementary Figure 6A).

## Discussion

### Summary

The current mechanistic hypothesis of amygdala neurofeedback is that the amygdala re-directs attention toward salient positive stimuli during self-referential processing, reducing rumination and improving forward-thinking [56, 57]. As observed in task-related and resting-state fMRI, these processes occur via the increased activation and functional connectivity changes in nodes comprising the salience and the default mode networks (SN and DMN, respectively) [9, 11, 36, 57, 58]. Cross-species neuroanatomical homologies [59, 60], including homologous SN [61], and DMN [62, 63] networks in the macaque brain, allow for a deeper delineation of these circuits involved in neurofeedback using NPHs. Previously, the NHP literature showed that the amygdala is anatomically interconnected with the large regions of the SN and DMN nodes [15, 16, 19, 20, 64]. Here, we provide tracer and dMRI evidence that the amygdala has monosynaptic anatomical connections with specific locations within the SN and DMN ipsilateral ROIs modulated by neurofeedback (Figure 6). We also show that amygdala hard-wiring with contralateral DMN ROIs is likely disynaptic through its connections with the adjacent hippocampus and PHG [14, 26, 28], two regions highly active during amygdala neurofeedback training [11, 36, 37]. This circuit delineation allows for new mechanistic descriptions of how the amygdala interactions with the SN and DMN could lead to lasting clinical effects after neurofeedback [8], as discussed below.

**Figure 6.**
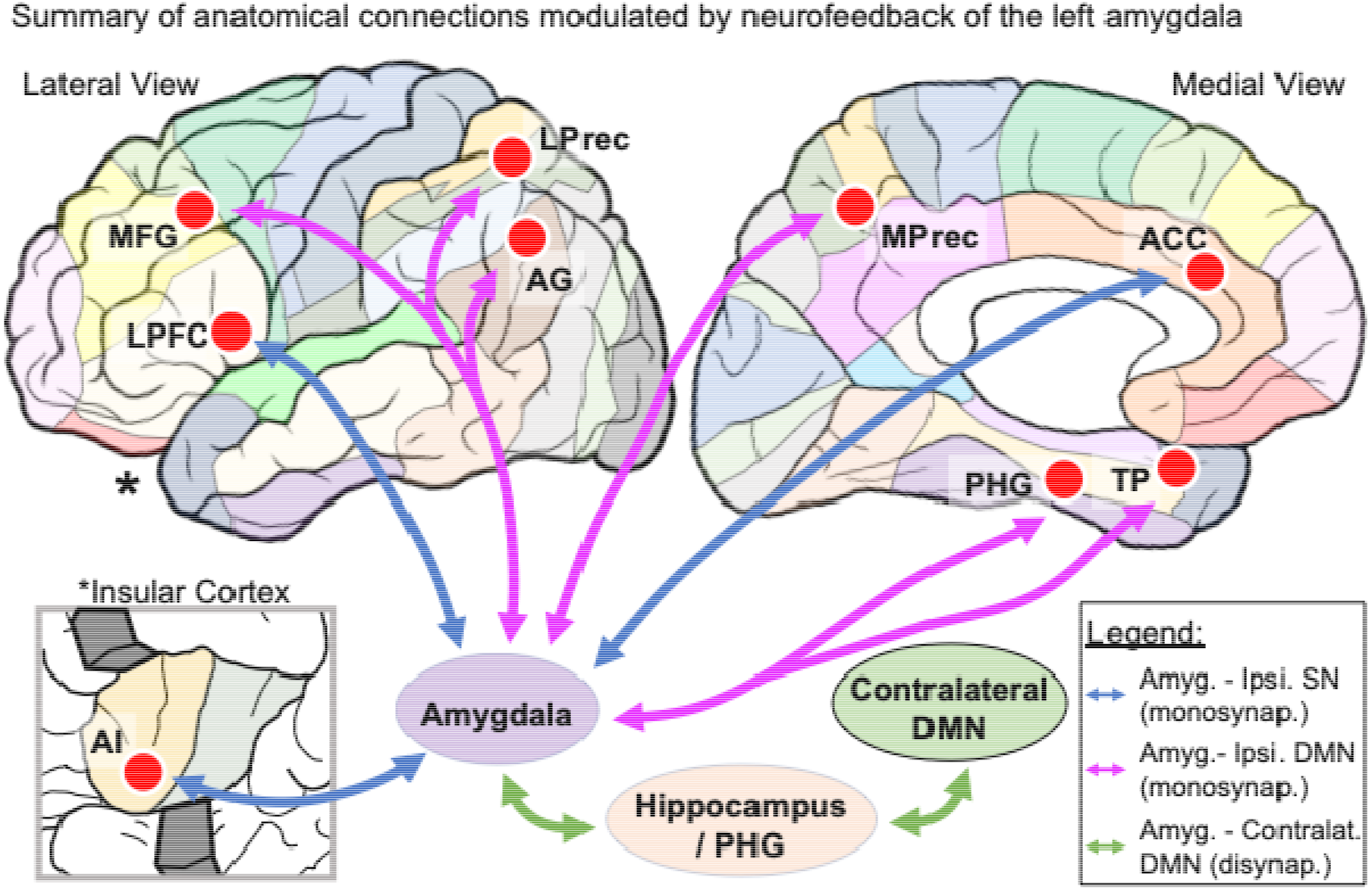
Summary of anatomical connections modulated by neurofeedback of the left amygdala. Representation of the specificity of ROIs in the left hemisphere (red circles) overlapping major functional regions (colored parcellation). Blue and pink arrows represent monosynaptic connections from the amygdala to the SN and DMN ipsilateral ROIs, respectively. Green arrows show the disynaptic connections with the DMN contralateral ROIs through the hippocampus and PHG.

### Amygdaloid connections to the ROIs within the SN and DMN

The amygdala connections to the SN nodes within the frontal and insular cortices are knowingly patchy and terminate in precise areas within each region [15, 19, 20, 22–24, 27, 29, 30]. Our results show that SN ROIs modulated by neurofeedback fall within these patches. These monosynaptic connections support the proposed role of amygdala neurofeedback re-directing attention toward specific salient stimuli [8, 65]. Previous NHP studies support that the amygdala works closely with the SN during salience processing [66–68]. E.g., local stimulation of the amygdala modulates the activity of the ACC and insular ROIs of the SN [69], reinforcing the potential of amygdala modulation of this network through its connections. Brain imaging and lesion studies in humans also highlighted the relevance of the amygdala and its connections in processing emotionally salient stimuli [70–74].

The amygdaloid connections with the DMN are less precise. For example, the amygdala has strong and widely distributed connections with the TP, Thalamus, Hippocampus, and PHG [13, 14, 17, 21, 22, 25, 26, 28–30]. However, connections with the MFG, PCC, and Precuneus are weaker and more restricted [15, 16, 18, 19, 29, 31]. Our data also showed that ipsilateral ROIs of the DMN modulated by neurofeedback are mainly connected with the central and basal nuclei. These nuclei are central for processing fear and anxiety [75–77], which are mediated by amygdala connections with regions processing context-specific aspects of the stress response [78, 79]. Modulation of fear and stress may play an essential role in worry and rumination, symptoms significantly correlated with amygdala rsFC with some DMN ROIs, including the MFG and precuneus [80].

Importantly, the ROIs listed in this study are specific to some but not all nodes of the SN and DMN. For example, regions like the ventral striatum (SN node) and vmPFC (DMN node) did not change rsFC with the amygdala after neurofeedback. These regions are known to be anatomically interconnected with the amygdala [15, 23, 24, 27, 29, 30, 32–34]. Although not identified in the rsFC studies, additional evidence suggests these regions are relevant during the neurofeedback task. The ventral striatum is highly active during neurofeedback reward processing [1]. Additionally, amygdala connectivity with the vmPFC changes during neurofeedback training [11, 81]. Thus, amygdala neurofeedback is associated with a modulation of the SN and DMN through anatomical connections.

Our data showed no monosynaptic connections from the amygdala to the contralateral ROIs, consistent with previous studies in the literature [82]. However, the amygdala is tightly linked with the ipsilateral hippocampus and PHG [14, 26, 28], which are connected to the contralateral structures [83–85]. Importantly, all studies included in our analysis [5, 12, 35] used a protocol based on positive autobiographical memory recall to up-regulate the BOLD signal of the left amygdala [11]. Neurofeedback studies using this protocol reported the hippocampus and PHG coactivation during the neurofeedback training task [11, 36, 37], increased functional connectivity between the left amygdala and left hippocampal/PHG structures [11, 12], and increased gray matter volume of hippocampal subfields [55]. Complementarily, neurofeedback targeting the up-regulation of the left hippocampus during autobiographical memory recall also leads to co-activation of the amygdala and increased amygdala-hippocampus functional connectivity [86]. Together with our anatomical delineation, these results suggest that rsFC changes with the contralateral DMN ROIs could be explained via amygdala-hippocampal projections.

### Neuroanatomical basis of clinical effects

The studies providing the ROI coordinates [5, 12, 35] are follow-up investigations from original trials with patients with depression [36, 87] or PTSD [88]. These patients showed significant clinical improvement and reduced symptoms after neurofeedback training [36, 87, 88]. Notably, around 30% of patients with depression reached remission levels at the primary endpoint [87]. These clinical effects correlated with the normalization of rsFC over the days following the neurofeedback training [5], similar to the effect observed in other protocols [6]. Thus, these clinical effects of fMRI neurofeedback training are likely to be associated with the rebalance of abnormal functional connections.

Monosynaptic connections allow the amygdala to modulate the ROIs of the SN and DMN quickly during neurofeedback sessions. A similar process to what is observed during focal stimulation of the amygdala [69, 89]: after systematic reinforcement, changes in these connections are sustained beyond the task and observed at rest [5, 12, 35]. These long-lasting connectivity changes may lead to synaptic rebalance and consequent clinical improvement, as observed in common pharmacological interventions [90].

### Technical considerations

Anatomical tract-tracing is the gold standard method for delineating connections in the primate brain [91]. However, our tracer data showed inconsistent labeling between the amygdala and lateral precuneus. In both cases, only retrograde labeling was identified. Proper tracer labeling in long-distance pathways may require up to 5 weeks of transport time [92, 93], while our cases were perfused after two weeks. Thus, a possible explanation is that the anterograde transport may need longer transport time to show labeling in long-distance connections. These transport characteristics should be considered in future studies.

For both species, we used submillimeter dMRI datasets (500 μm in NHPs and 760 μm in humans) to delineate bundles that would be inaccessible at lower-resolution [94]. However, even at the submillimeter scale, the reconstruction of some anatomical connections identified in the tracer data was challenging in the dMRI data. For example, very few streamlines were identified linking the amygdala and the LPFC in both species. However, NHP tracer data show amygdala projections traveling through the uncinate fasciculus to reach their targets in the ventrolateral prefrontal cortex [95, 96], with similar fiber organization in the human brain [97]. In both species, dMRI data also showed false positive connections with the contralateral hemisphere. Some of these contralateral connections identified using dMRI are likely caused by the proximity of the fornix to actual amygdala pathways, such as the stria terminalis (<700 μm). In fact, studies trying to separate these bundles also reported the partial volume effects in their tract reconstructions [98, 99]. Therefore, the combined analysis of NHP tract tracing and NHP and human dMRI data is essential for delineating circuits relevant to neuroimaging studies [91] and identifying challenging fiber configurations for tractography algorithms [100–102].

### Conclusion and Future Perspectives

We described the neuroanatomical circuits involved in rsFC changes following amygdala neurofeedback training. We showed that neurofeedback modulates the SN and DMN through monosynaptic connections from the targeted region providing real-time feedback (amygdala) and disynaptic connections with areas involved in the targeted cognitive process (hippocampus and PHG during autobiographical memory recall). Such circuitry allows for rapid modulation and reinforcement of amygdala connections with large-scale networks, leading to clinical improvements observed in the literature. This new mechanistic hypothesis should be probed in future human and animal studies. Moreover, our approach (combining NHP tract-tracing and dMRI *ex vivo* and human dMRI *in vivo*) can also guide future clinical neurofeedback experiments. For instance, it allows the identification of amygdala nuclei for targeting with neurofeedback at high field (e.g., as those achievable at 7T). It can also select alternative targets within the anatomical circuit to optimize neurofeedback for non-responders to the amygdala modulation.

## Acknowledgments

LT was supported by the Jonathan Edward Brooking Mental Health Research Fellowship and NIH grant K99-MH130648. LT, JL, and SH were partially supported by NIH grants P50-MH106435 and R01-MH045573. CM, ED, and AY were partially supported by NIH grants R01-NS119911 and R01-EB021265.

