## SUPPLEMENTARY INFORMATION for "Translation of monosynaptic circuits underlying amygdala fMRI neurofeedback training"

### Methods

*Anatomical injections*

The University Committee on Animal Resources from the University of Rochester approved all tracer experiments, and animal care followed the National Guide for the Care and Use of Laboratory Animals.

We identified the stereotaxic coordinates for the injection sites using pre-surgery structural MR images. Monkeys received injections of one or more of the following tracers: Lucifer Yellow (LY), Fluororuby (FR), Fluorescein (FS) (40 –50 nl, 10% in 0.1 M phosphate buffer [PB], pH 7.4; Invitrogen), or tritiated amino acids (100 nl, 1:1 solution of [3 H] leucine and [3 H]-proline in dH2O, 200 mCi/ml, NEN). Tracers were pressure-injected over 10 min using a 0.5 l Hamilton syringe. After each injection, the syringe remained *in situ* for 20-30 min.

Twelve to 14 days after the surgery, monkeys were deeply anesthetized and perfused with saline, followed by a 4% paraformaldehyde/1.5% sucrose solution. We post-fixed brains overnight and cryoprotected in increasing sucrose gradients [1]. We cut serial sections of 50 mm on a freezing microtome, and processed one in every eight free-floating sections to visualize LY, FR, FS, and AA tracers, as previously described [2, 3]. We mounted sections onto gel-coated slides, dehydrated, defatted in xylene overnight, and coverslipped with Permount. In cases with more than one tracer injection into a single animal, we processed adjacent sections for each antibody reaction.

*NHP dMRI acquisition*

NHP samples were packed in a bag, filled with fomblin, and occasionally manually massaged over two days to remove air bubbles trapped within the brain. We left the samples out at room temperature for a minimum of 6 hours prior to scanning. Samples were scanned in a small-bore 4.7T Bruker BioSpin MRI system, with a gradient internal diameter of 114 mm, maximum gradient strength 660 m/Tm, and a birdcage volume RF coil internal diameter of 72 mm. A two-shot 3D echo-planar imaging (EPI) sequence was used for dMRI with TR = 500 ms, TE = 48 ms, $\delta$ = 15 ms, $\Delta$ = 19 ms, $b_{max}$ = 40,000 s/mm$^2$, 514 gradient directions, and 0.5 mm isotropic resolution. The above scanning parameters result in a total scan time of 47 hours for each NHP sample.

*Human dMRI acquisition*

We used high-resolution diffusion MRI (dMRI) data from a publicly available and pre-processed dataset [4]. Briefly, data were acquired on the MGH-USC 3T Connectome Scanner at 0.76 mm isotropic resolution using an SNR-efficient simultaneous multi-slab imaging technique (gSlider-SMS) [5, 6] across 9 2-hour sessions (max gradient amplitude: 180 mT/m; slew rate: 125 T/m/s, gSlider factor: 5, MB factor: 2, R: 3, TR/TE: 3500/75 ms, matrix: 290x288, PE: AP). 2808 dMRI volumes were acquired (144 b=0, 420 b=1000, 840 b=2500 s/mm$^2$, and their paired reversed PE volumes).

*Cross-species brain parcellation*

The "Regional Map" (RM) parcellation was initially manually drawn on the F99 macaque brain template [7]. Later, it was deformed to surface- and volume-based human MNI representations [7, 8] using a landmark-based deformation defined by a set of major 150 sulci and gyri, along with functional activation patterns, considered homologous between the two species [9].

### Results

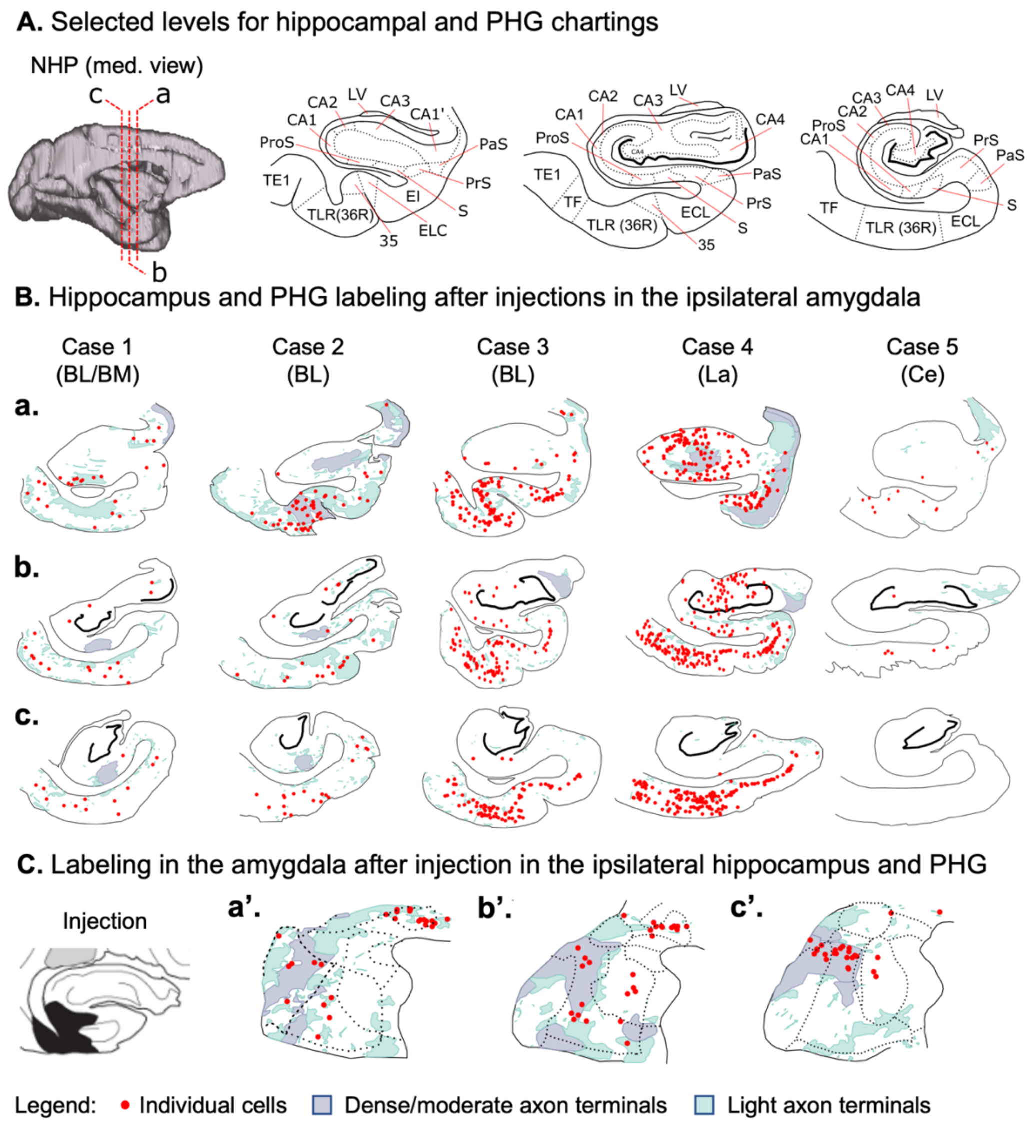


**Supplementary Figure 1 – Amygdala connections with the hippocampus and parahippocampal gyrus (PHG).** (**A**). 3D representation of the three rostro-caudal levels (a-b) used in the chart cells and terminals in the hippocampus and PHG, and the respective coronal slices with cytoarchitectonic divisions based on the Paxinos atlas [10]. Labeling of cells (red dots), and dense/moderate (light blue) and diffuse (light green) terminal fields in the hippocampus and PHG after bidirectional tracer injections in different amygdala nuclei (**B**). In (**C**), the labeling of cells and terminals in the amygdala (following the coronal levels a'-c' from Figure 5) after a bidirectional tracer injection in the hippocampus and PHG (left). *Abbreviations:* 35 = area 35 of cortex, BL = basolateral nucleus, BM = basomedial nucleus, C = caudal, CA1 = field CA1 of the hippocampus, CA1' = field CA1' of the hippocampus, CA2 = field CA2 of the hippocampus, CA3 = field CA3 of the hippocampus, CA4 = field CA4 of the hippocampus, Ce = central nucleus, ECL = caudal limit part of the entorhinal cortex, EI = intermediate part of the entorhinal cortex, ELC = lateral part of the entorhinal cortex, La = lateral nucleus, LV = lateral ventricle, PaS = parasubiculum, ProS = prosubiculum, PrS = presubiculum, R = rostral, S = subiculum, TE1 = temporal area TE1, TF = temporal area TF, TLR(36R) = rostral part of the temporal area TL (area 36R).

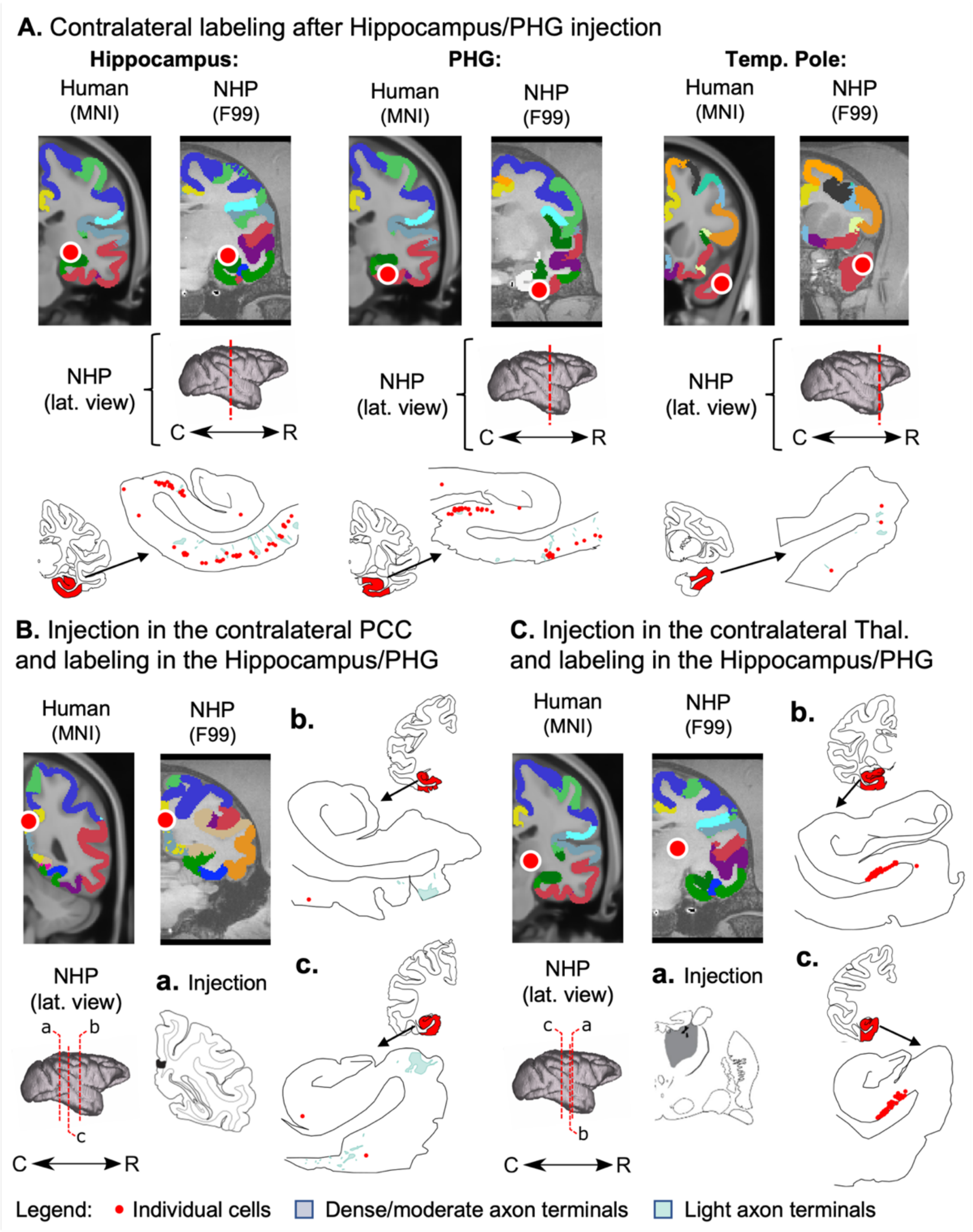


**Supplementary Figure 2 – Hippocampus and parahippocampal gyrus (PHG) connections with contralateral DMN nodes.** Red circles indicate the peak location of the rsFC changes after amygdala neurofeedback for the right Hippocampus (**A** left), PHG (**A** center), and Temporal Pole (**A** right). 3D models represent the rostro-caudal level from each node. For each region, schematic coronal sections highlight in red the location with connectivity chartings zoomed in showing labeling of cells (red dots) and dense/moderate (light blue) and diffuse (light green) terminal fields after bidirectional tracer injections in the contralateral Hippocampus and PHG (injection shown in Supplementary Figure 1C left). ROIs in the right PCC (**B**) and Thalamus (**C**) and respective injections also show labeled cells and terminals in the contralateral hippocampus and PHG. *Abbreviations:* BL = basolateral nucleus, BM = basomedial nucleus, C = caudal, Ce = central nucleus, La = lateral nucleus, R = rostral.

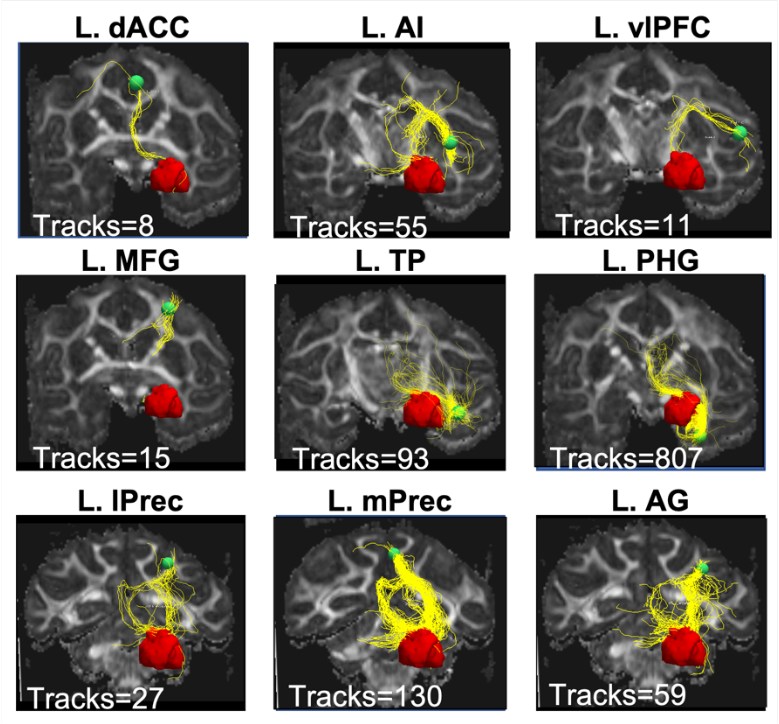


**Supplementary Figure 3 – dMRI tractography results for animal 2.** Examples of amygdala seeds (red) and the tracts (yellow) connecting them with all ipsilateral nodes (green) within the SN and DMN.

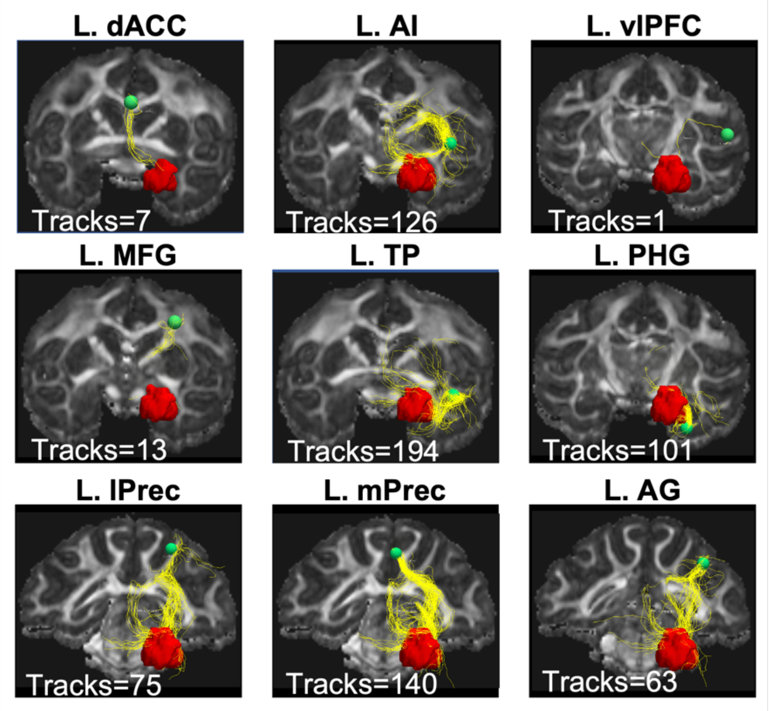


**Supplementary Figure 4 – dMRI tractography results for animal 3.** Examples of streamlines (yellow) connecting the amygdala (red) with all ipsilateral nodes (green) within the SN and DMN.

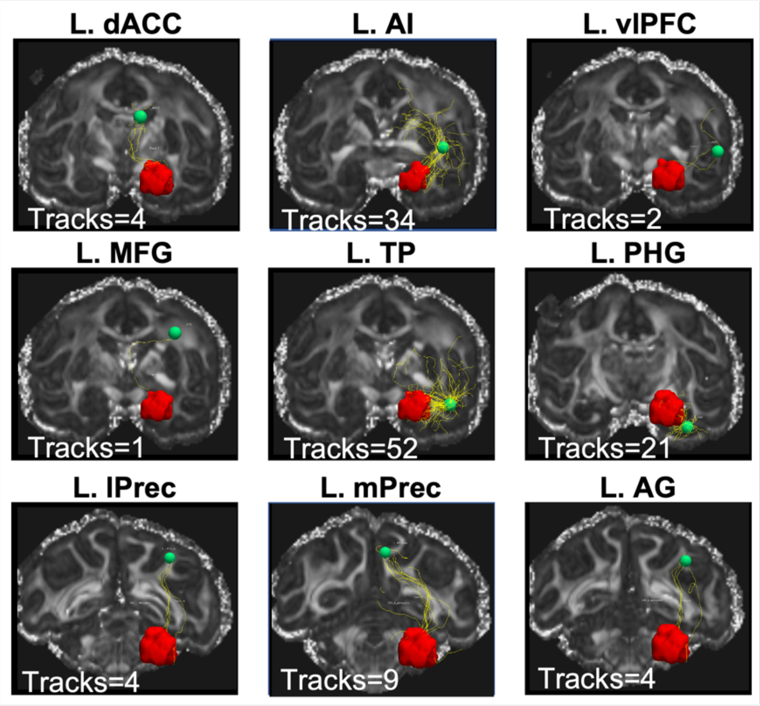


**Supplementary Figure 5 – dMRI tractography results for animal 4.** Examples of amygdala seeds (red) and the tracts (yellow) connecting them with all ipsilateral nodes (green) within the SN and DMN.

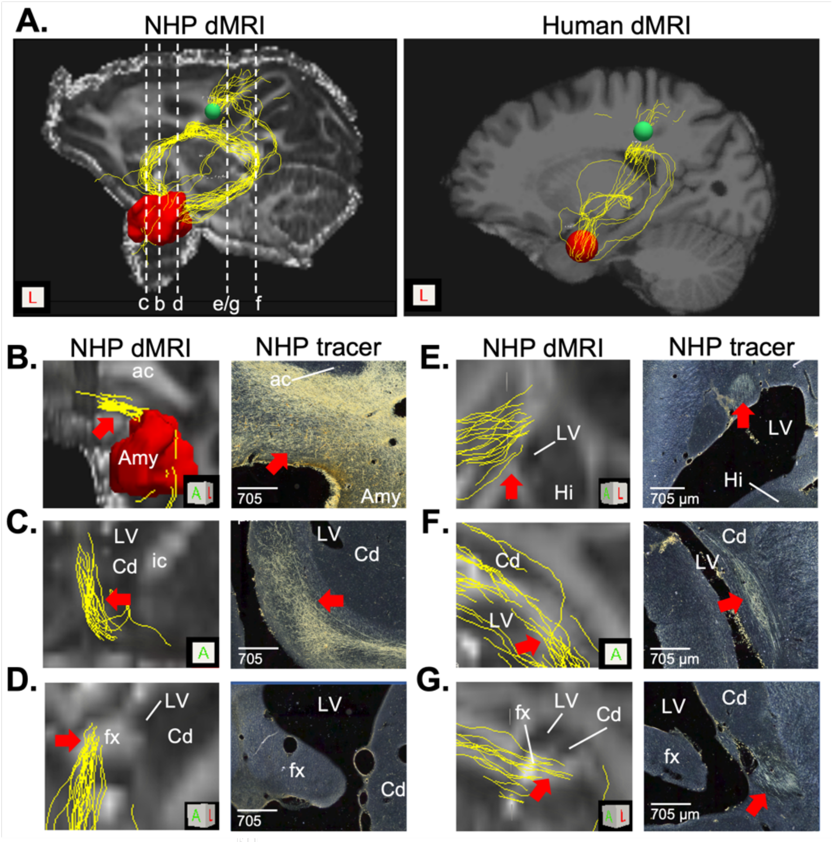


**Supplementary Figure 6 – Tract tracing data allows for identifying false positive tractography results.** (**A**) Results in the NHP and human brains show two major tracts (anterior and posterior) connecting the left amygdala and right medial precuneus. The most anterior tract follows the fibers of the amygdalofugal pathway from the amygdala (**B**) to the medial wall (**C**) until they are captured by the fornix (**D**). The posterior tract follows fibers from the stria terminalis (E-G) until they erroneously follow the fornix streamlines. *Abbreviations:* ac = anterior commissure, Amy = amygdala, cc = corpus callosum, Cd = caudate nucleus, fx = fornix, Hi = Hippocampus, ic = internal capsule, LV = lateral ventricle.
